# Population genetic structure of the aphid pest *Myzus persicae* in organic and conventional greenhouses

**DOI:** 10.1101/2024.07.22.604537

**Authors:** Mariska M. Beekman, Joost van den Heuvel, S. Helena Donner, Jordy J. H. Litjens, Marcel Dicke, Bas J. Zwaan, Eveline C. Verhulst, Bart A. Pannebakker

## Abstract

Aphids display remarkable adaptability to pest control strategies and their parthenogenetic reproduction results in rapid numerical increase of higher-fitness clones. Consequently, a high prevalence of only a few clonal lines could indicate positive selection for these genotypes, potentially reflecting adaptation to pest control methods. Here, we investigated the clonal diversity and population genetic structure of the green peach aphid *Myzus persicae* using microsatellite markers, in both organic and conventionally managed Dutch sweet pepper greenhouses over four consecutive years. In total, 26 distinct multilocus genotypes (MLG) were detected, with higher clonal diversity in organic than in conventional greenhouses. Strikingly, a single MLG dominated conventional greenhouses in 2019, only to be completely replaced by a new dominant MLG by 2022. Whole-genome sequencing of 15 sampled lines — seven sharing the same MLG and eighth with unique MLGs — revealed that all aphids with the dominant MLG of 2019, collected from various locations and over multiple years, originated from a single parthenogenetic ancestor. Our findings indicate that the population genetic structure of *M. persicae* differs between organically and conventionally managed sweet pepper greenhouses. The presence of dominant MLGs in conventional crop systems may suggest positive selection or evolutionary forces such as a founder effect. Understanding the forces driving these differences in population genetic structure and their impact on the efficacy of biocontrol agents will further help us improve control strategies for *M. persicae* in greenhouse crops.

## Introduction

Insect herbivore populations often evolve remarkably rapidly under the strong and distinct selection pressures of pest control strategies applied in agricultural systems (Chen and Schoville, 2018; Pélissié et al., 2018). Agricultural systems, and especially commercial greenhouses where the biotic and abiotic environment are strictly controlled, are often much more homogeneous and simplified than natural environments. These characteristics are linked with lower levels of natural pest control and increased pest pressure (Bianchi et al., 2006; Rusch et al., 2016). Consequently, commercial greenhouses are especially susceptible to herbivore pest populations. Control of pest populations in these systems is vulnerable to the evolution of and selection for resistance to various pest management strategies, including chemical insecticides (Mota-Sanchez and Wise, 2023), bio-insecticides (Siegwart et al., 2015), insect-resistant transgenic crops (Tabashnik and Carrière, 2015), or natural enemies used as biocontrol agents (Tomasetto et al., 2017). Early detection of pest populations adapting to control strategies is thus crucial for effective pest management in commercial greenhouses.

Aphids are highly damaging agricultural insect herbivores (Dedryver et al., 2010) with diverse mechanisms for withstanding control strategies. For example, the use of chemical insecticides, historically the primary means of aphid control, have led to multiple aphid species developing resistance to various aphicidal compounds (Mota-Sanchez and Wise, 2023). In addition, aphids exhibit ample variation in their defensive behaviours towards predators (Braendle and Weisser, 2001; Dion et al., 2011; Kunert et al., 2010), and in their defence mechanisms against parasitoid wasps and entomopathogenic fungi, which are all used in biocontrol. These defence mechanisms consist of 1) genetically encoded defences, also known as endogenous resistance (e.g. Martinez et al., 2014; Parker et al., 2014; Sandrock et al., 2010; Von Burg et al., 2008), and 2) symbioses with protective facultative endosymbionts that are maternally transmitted to the offspring (e.g. Łukasik et al., 2013; Oliver et al., 2003). Therefore, when biocontrol agents are deployed to control aphid populations, the resulting selection force can act on traits encoded by the complete heritable aphid hologenome (meaning the collective genomes of the host and its microbiome; Bordenstein and Theis, 2015).

Aphids are exceptionally proficient agricultural pests due to their distinct reproductive mode, known as cyclical parthenogenesis or a holocyclic life cycle (Moran, 1992). In cyclical parthenogens, new genotypes are formed during a single yearly sexual cycle, which is triggered by environmental changes such as colder temperatures and shorter days (Lees, 1966). The rest of the year, aphids reproduce by apomictic parthenogenetic (asexual) reproduction, resulting in clone-mates (individuals originating from a single parthenogenetic ancestor) that share the same genotype. However, aphid lineages can also exhibit an anholocyclic life cycle, reproducing parthenogenetically year-round, either due to genetic or environmental changes. For instance, aphid lineages can develop into obligate parthenogenetic lineages, completely losing the ability for sexual reproduction. Alternatively, lineages can skip sexual reproduction while still being able to. This occurs when environmental cues or primary hosts required to trigger sexual reproduction are absent, as observed in the homogeneous and sheltered environment of commercial greenhouses. In anholocyclic lineages, favourable gene combinations are no longer broken up during meiosis, allowing for the selection and expansion of highly successful genotypes, which may be maintained over large geographical areas for an extended period of time. Consequently, a lack of clonal diversity, or the dominance of specific clones (clonal unevenness) in a greenhouse crop may suggest selection and adaptation in aphid populations.

Of all aphid species, the green peach aphid, *Myzus persicae* (Sulzer), has the highest worldwide economic significance, annually resulting in substantial yield losses and tremendous costs associated with its control. It has a cosmopolitan distribution (Margaritopoulos et al., 2009), can transmit many different plant viruses (Nault, 1997; van Emden and Harrington, 2017), is well-known for its remarkable ability to quickly develop resistance to many insecticides (reviewed by Bass and Nauen, 2023; Bass et al., 2014), and has a broad host range including many agriculturally important crops (Blackman and Eastop, 2000). *Myzus persicae* reproduces sexually only on *Prunus* ssp. Therefore, the influx of new clones of *M. persicae* into a greenhouse system, where *Prunus* ssp. are absent, is expected to be much lower than in natural systems as new clones can only enter the greenhouse physically and not through sexual reproduction.

In this study, we focused on *M. persicae* as a problematic pest in commercial Dutch sweet pepper greenhouses. Effective control, both in conventionally (chemical insecticides and biocontrol agents are both deployed) and organically (excluding all input from chemical sources) managed systems, is often difficult to achieve (Glastuinbouw Nederland, 2022; Messelink et al., 2011). Although the reasons for these difficulties remain unclear, they were suggested to include protection by facultative symbionts and endogenous resistance (Beekman et al., 2022; Vorburger, 2018). However, our previous work showed that *M. persicae* from Dutch sweet pepper greenhouses does not carry facultative endosymbionts (Beekman et al., 2022). In the same study, we also did not observe any variation in endogenous resistance to the biocontrol parasitoids *Aphidius colemani* Vierick and *Aphidius matricariae* Haliday between various clonal lines of *M. persicae* collected from Dutch sweet pepper.

To unravel why *M. persicae* is such a problematic pest species in sweet pepper, we investigated the clonal diversity and population genetic structure of *M. persicae* in Dutch sweet pepper crops over four successive seasons. We aimed to determine whether these populations are multiclonal, or if specific clonal lines dominate or become dominant. As distinct pest control methods are applied in conventionally versus organically managed greenhouses, we hypothesise that different genotypes thrive under each management type. Therefore, we analysed the clonal diversity and prevalence of specific clones for *M. persicae* population from both types of Dutch sweet pepper greenhouses separately.

## Materials and methods

### Aphid collection

*Myzus persicae* samples were collected from sweet pepper, *Capsicum annuum* L. (Solanaceae), greenhouses (enclosed glass buildings used for commercial crop production) located in the Netherlands. We sampled conventional greenhouses (using both chemical insecticides and biocontrol agents for pest control) during various time periods during the crop growth season from 2019 up until 2022, and organic greenhouses (excluding all chemical pesticides, only using biocontrol agents) in 2019 and 2022 (Figure 1). The sweet pepper growth season in the Netherlands roughly starts in January and runs until December. In total, aphids were sampled during nine different time periods; three in 2019, and two in 2020, 2021, and 2022. Details on greenhouse locations, sampling dates, and number of genotyped aphids, are presented in Suppl. Table S1.

**Figure 1.**
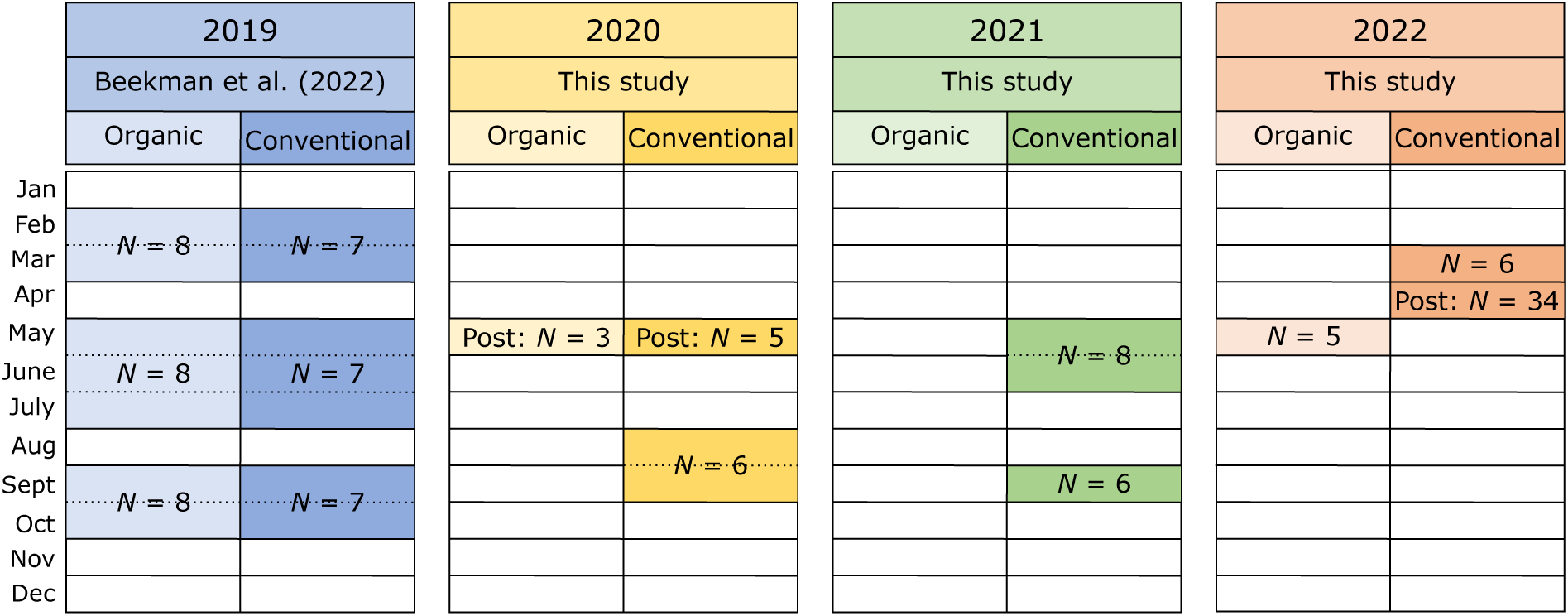
Overview of *Myzus persicae* sampling in Dutch sweet pepper greenhouses and how many greenhouses were sampled (“*N*”). The 2019 samples were collected as part of a previously published study (Beekman et al., 2022), while all other samples were collected as part of the current study. “Post” represents samples that were sampled by growers and sent to us by post instead, all other sampling was performed by the researchers.

Aphids were collected either as single individuals or, whenever possible, together with other aphids from the same colony present on a single leaf and stored in 70% EtOH. Aphids were morphologically identified as *M. persicae* using the identification keys of Blackman and Eastop (2000), after which the samples were stored at -20 °C until further use.

### Sampling by the researchers 2019-2022

During every sampling time period in 2019, samples from eight conventional and seven organic greenhouses were obtained (Figure 1). The 2019 samples, collected as part of a different study (Beekman et al., 2022), were collected by dividing greenhouse compartments into 6 x 6 grids. Each cell of the grid was partially sampled and when aphids were detected, multiple aphids of a single colony were collected. From 2020 onwards, the sampling strategy was modified, and greenhouse compartments were sampled less thoroughly. Aphids were only searched for in the four outermost corners of the greenhouse, as well as in two to four additional spots randomly picked in between the four outermost corners, when applicable at hotspots identified by the growers. In August-September 2020, five conventional greenhouses were sampled. In May-June 2021 and September 2021, we sampled eight and six conventional greenhouses respectively, and in 2022, six conventional and five organic greenhouses were sampled in March and May respectively.

### Additional sampling by the growers in 2020 and 2022

In May 2020, because of COVID-19 restrictions, sampling by researchers was not possible and growers of previously sampled greenhouses were asked to collect and send in a single sample per greenhouse. In this way, samples from five conventional and three organic greenhouses were received (Figure 1).

To enable monitoring of specific genotypes across the Netherlands, additional Dutch sweet pepper growers were invited to voluntarily participate in a large-scale sampling effort of *M. persicae* from their greenhouses in April 2022 (see Suppl. Table S2 for information about the greenhouses). Participating growers were asked to sample up to six different aphid colonies from various hotspots, as far away from each other as possible, in their greenhouse. Furthermore, they were asked to fill out a questionnaire about the farming management type (conventional or organic), the growing conditions, perceived difficulties with aphid infestations, and, when applicable, which aphid-targeting insecticides they had applied in their greenhouses (see Suppl. File S1 for the original questionnaire). Care was taken to instruct the growers to follow the same sampling strategy as the researchers, but possible sampling biases cannot be ruled out. Additionally, the voluntary nature of grower participation in this study may have introduced a self-selection bias, potentially favouring the inclusion of growers facing more challenging aphid genotypes. Sampling by the growers yielded an additional 194 samples from 34 conventional greenhouses in April 2022.

### Aphid lines

Living *M. persicae* aphids were sampled in 2019, 2021 and 2022 (Table 2) to establish laboratory lines. Lines were started from single parthenogenetic females and were reared in sterile polypropylene culture vessels (Lab Associates, Oudenbosch, the Netherlands) closed with nylon-screened Donut Lids (BDC0001-1; Bugdorm, MegaView Science, Taichung, Taiwan), on sweet pepper (*Capsicum annuum* L.) leaf discs placed on 1% agar, stored upside down at 15 °C, 16-hour light:8-hour dark photoperiod, and 60% relative humidity in an incubator (Sanyo MLR-352H, PHC Europe, Etten-Leur, The Netherlands). Aphid lines were named after their multilocus genotype (MLG), followed by the identifier of the greenhouse (see Suppl. Table S1), and the year they were collected in.

### Genotyping through microsatellite analysis

Aphids were genotyped using multi-allelic microsatellite markers, which are useful to distinguish clonal lines in parthenogenetic organisms such as aphids. We targeted the previously published microsatellite loci M37, M40, M86 (Sloane et al., 2001), myz2, and myz9 (Wilson et al., 2004). From each greenhouse compartment, a maximum of four aphids located the furthest apart from each other were chosen for genotyping. This selection was made under the assumption that if these aphids shared the same MLG, it would be likely that the entire greenhouses harboured a single MLG. In instances where different *M. persicae* colour morphs were observed in a greenhouse compartment, the above-described principle was applied for both colour morphs. When multiple MLGs were identified in a greenhouse compartment, additional genotyping was conducted on samples collected in between the identified MLGs. This approach aimed to comprehensively capture the genetic diversity within the greenhouse during each sampling time period. For the exact number of aphids genotyped during each time period, see Suppl. Table S1.

DNA was extracted from the EtOH-stored aphids, which were dried on tissue paper before homogenizing whole individuals in 100 µL of 5% Chelex-100 resin (Bio-Rad, Hercules, CA, USA) in ultrapure water with 2.5 µL proteinase K (20 mg/mL; Promega, Southampton, UK). The samples were incubated for a minimum of 60 minutes at 56 °C, followed by heating to 98 °C for 8 minutes to deactivate the proteinase K. The resulting supernatant, containing DNA, was diluted at 1/3 in ultrapure water and subsequently stored at -20 °C until being used for PCR amplification.

In the PCR assays, all forward primers were fluorophore-tagged using universal tails, following the ‘Protocol for concurrent primer labelling and multiplexing’ by Culley et al. (2013) and using the universal tails ‘A’ and ‘B’ by Blacket et al. (2012). This cost-efficient method entails indirect fluorescent labelling of forward primers by introducing a fluorescently labelled universal primer corresponding to a universal tail added to the forward primers, in the PCR mix. Locus-specific forward primers M86 and myz9 were tailed with universal tail ‘A’ whilst M37, M40 and myz2 were tailed with universal tail ‘B’. Universal primer ‘A’ was tagged with fluorophore PET and universal primer ‘B’ with 6-FAM.

PCR reactions were carried out in 5 µL-volumes, comprising 1x reaction buffer QIAGEN Multiplex PCR Master Mix (containing dNTPs, HotStartTaq DNA polymerase and 3 mM MgCl_2_ as final concentration; QIAGEN, Venlo, the Netherlands), 0.05 µM tailed forward primer (see Suppl. Table S3 for all primers), 0.2 µM reverse primer, 0.2 µM fluorophore labelled universal primer, and 1 µL DNA. Multiplexing was employed for markers M37 and myz2 (multiplex mix 1), as well as for M40 and M86 (multiplex mix 2), while myz9 (multiplex mix 3) was amplified individually due to the formation of primer dimers when combined with other primers. Thermocycler conditions were as follows: initial denaturation at 95 °C for 15 min; 30 cycles of 94 °C for 30 s, 57 °C for 45 s, and 72 °C for 45 s; 8 cycles of 94 °C for 30 s, 53 °C for 45 s, and 72 °C for 45 s; followed by 72 °C for 20 min.

Following PCR amplification, the three PCR products per aphid sample were combined in equivolume ratios for subsequent fragment analysis. Fragment separation was carried out using a 3730XL DNA Analyzer (Applied Biosystems, Waltham, Massachusetts, USA). Electropherograms were visualized, and allele sizes were determined using Geneious Prime v. 2019.1.3 (BioMatters Ltd., Auckland, New Zealand) with the Microsatellite Plugin v. 1.4.7. Samples lacking data for any of the markers were omitted from further analyses.

The assignment of names to MLGs involved designating unique identifiers to those differing at multiple loci or exhibiting variation of multiple repeats at a single locus, denoted as ‘MLG-A’, ‘MLG-B’, ‘MLG-C’, etc. Multilocus genotypes closely resembling more prevalent MLGs, differing by only a single repeat at a single locus, and discovered in the same greenhouse and at the same sampling point as their more common near-identical MLG (henceforth called ‘sister MLG’) were considered near-identical by descent and named ‘MLG-A2’, ‘MLG-A3’, ‘MLG-B2’, etc. However, as this near-identicality can also be due to homoplasy, near-identical MLGs not detected together with their sister MLG received unique names, similar to all other unique MLGs.

### Data analyses

Data analyses were performed in R v. 4.2.1 (R Core Team, 2022) using RStudio v. 22.07.0 (RStudio Team, 2020). The R package *poppr* v. 2.9.4 (Kamvar et al., 2014, 2015), designed for genetic analyses analysis of partially clonal populations, was used to analyse the microsatellite data.

Multilocus genotypes were identified for all samples using the *mll* function, genotypic richness with the *poppr* function, and allelic richness with *locus_table* function. All following indices were calculated on all data, excluding the samples collected by large-scale sampling of the growers themselves in 2022.

As we sampled *M. persicae* from sweet pepper, a secondary host on which it reproduces parthenogenetically, deviations from Hardy-Weinberg equilibrium (HWE) and increased linkage among the microsatellite markers are expected. We analysed the markers for Hardy-Weinberg equilibrium (HWE) using the *hw.test* function (chi-square goodness-of-fit test) of the R package *pegas* v. 1.3 (Paradis, 2010) with 1000 permutations, per sample group (organic and conventional) as well as for all samples together. To test for linkage among markers per population, we calculated the index 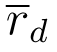 (Agapow and Burt, 2001) which is based on the index of association *I_A_*(Brown et al., 1980), but accounts for the number of loci sampled, using the *ia* function with 1000 permutations on both the normal and clone-corrected data. 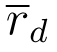is expected to be zero in a freely recombining population, and larger than zero when alleles are associated. Observed heterozygosity (*H_o_*) was calculated using the *summary* function.

Visual representations of data were computed with the *ggplot2* package v. 3.4.4 (Wickham, 2016) and modified using Inkscape v. 1.2.2 (Inkscape Project, 2022).

### Whole genome resequencing

To assess whether five microsatellite markers were sufficient to produce unique MLGs that represent aphids that all descended from the same parthenogenetic ancestor and are thus true clone-mates, we sequenced the whole genomes of 15 *M. persicae* lines. These aphid lines were established as laboratory lines as described above under section ‘Aphid lines’. Each line originated from a distinct greenhouse and/or unique sampling time period, representing nine MLGs. Seven of the sequenced aphid lines shared the same MLG, which dominated in 2019. The remaining eight lines all had distinct and unique MLGs.

DNA was extracted from approximately 15 adult aphids per line, using the DNeasy Blood & Tissue Kit (QIAGEN) following the manufacturer’s instructions. DNA was eluted in RNAse-free water and treated with RNAse ONETM (Promega) before being sequenced on an Illumina NovaSeq, generating 150-bp paired-end reads, in two separate sequence runs. The first run included nine samples collected in 2019 and the second run included six samples collected in 2021.

The data was analysed using the following steps. First, raw reads were quality filtered with Trimmomatic v. 0.33 (Bolger et al., 2014) using default settings, and size filtered for a minimum length of 70 bp. To confirm the identity of the sequenced aphid lines as *M. persicae*, the obtained sequence reads were aligned with the *M. persicae* mitochondrial genome (Jung et al., 2021) using Geneious Prime (BioMatters Ltd.). The alignments with the cytochrome c oxidase subunit I (COI) region were checked against the Barcode of Life Data System (BOLD; available at https://www.boldsystems.org/) for correct species identification.

Subsequently, the quality- and size-filtered reads were aligned with the *M. persicae* reference genome of clone G006 v. 3 (available at https://bipaa.genouest.org/is/aphidbase/) using BWA v. 0.7.17 (Li, 2013) and SAMtools v. 1.14 (Li et al., 2009). The alignment quality was assessed using SAMtools flagstat, and duplicate reads were removed with PICARD’s MarkDuplicates tool v. 2.8.3 (http://broadinstitute.github.io/picard/). Variants were called using FreeBayes v. 0.9.21 (Garrison and Marth, 2012), resulting in a VCF file, which was filtered with bcftools v. 1.10.2 (Danecek et al., 2021) for a minimum depth of 10 and a minimum quality of 20.

### Identification of repeat regions

Repeats were identified using the SNPable Regions pipeline (http://lh3lh3.users. sourceforge.net/snpable.shtml). The reference genome was split into 50 basepair k-mers using the *splitfa* function, which were mapped to the reference genome using BWA-aln and BWA samse under default settings (Li, 2013). A masked FASTA file was produced using *gen_mask* (-l 35 -r 0.5), and a custom Perl script was used to produce a BED file from the lower case bases, which represent repeat regions, from the masked FASTA file.

### Genetic diversity estimates

Nucleotide diversity, Waterson’s estimator and Tajima’s D were estimated using PoPoolation1 v. 1.2.2 (Kofler et al., 2011). First, pileup files were generated for each sample, from the deduplicated BAM files, using SAMtools *mpileup*. To ensure equal contribution for each file, the pileup files were subsampled using the subsample-pileup.pl script from PoPoolation1 with a minimum coverage of 10, a maximum coverage of 80, and a minimum quality of 20, without replacement (Kofler et al., 2011). Subsequently, the PoPoolation1 filterpileup-by-gtf.pl script was used to transform the repeat regions BED file into a GTF file, to filter out repeat regions from the pileup files. All pileup files were combined to produce a unified pileup for all samples. The coalescence measures were estimated using a sliding window approach with the PoPoolation1 script Variance-sliding.pl with –window-size 500000 –step-size 250000 –min-count 2 –pool-size 500 –min-coverage 140 –max-coverage 160 –min-qual 0 –fastq-type sanger and –min-covered-fraction 0.1.

The previously produced VCF file was filtered for repeat regions using the BEDtools v. 2.27.1 tool intersect (Quinlan and Hall, 2010). Next, a custom R script was employed to filter variants based to the following criteria: minor allele counts over all samples had to exceed 2%, the mean coverage had to be higher than 2.5% and lower than twice the median of this coverage distribution, variants should be biallelic, the distance to indels should be greater than nine basepairs, and multi-nucleotide variants were restricted to a size of four basepairs. Additionally, only variants with genotype calls and read coverage *≥* 10 in all 15 samples were retained. The resulting filtered VCF was then used to calculate heterozygosity using the *het* function of VCFtools v. 0.1.13 (Danecek et al., 2011), by subtracting the number of observed and expected homozygous sites from the total number of sites.

Genetic distances were estimated by assigning a value of 0 to reference homozygotes (“0/0”), a value of 1 to alternative homozygotes (“1/1”), and a value of 0.5 to heterozygotes. This approach, based on the concept of allele frequency distance (Berner, 2019), encoded the genetic distance per locus between two alternative homozygotes as 1, and between a homozygote and a heterozygote as 0.5. However, we did not divide the variant number of loci by the total genome size, but counted the total allele frequency distance. A distance of 100,000 therefore would translate into 100,000 loci that were homozygous reference for one aphid line and homozygous alternative for a second aphid line. We used hierarchical clustering using this measure as a distance measure to cluster samples with the use of the R base function *hclust* with ‘average’ as the agglomeration method. Lastly, we counted unique shared mutations among aphid lines, focusing on shared heterozygous sites that were homozygous in one direction, either “0/0” or “1/1”, for all other aphid lines.

## Results

### Population genetic structure of *Myzus persicae* in sweet pepper

#### Multilocus genotypes and microsatellite diversity

A total of 348 colonies of *M. persicae* were sampled by the researchers over 2019-2022, 204 from conventional and 144 from organic greenhouses, and an additional 194 samples were sent in by growers in 2022. All samples were genotyped at five microsatellite loci and a total of 26 MLGs were detected, which were called ‘MLG-A’, ‘MLG-B’, ‘MLG-C’ etc. (Figure 2; Suppl. Table S4). Twenty-two of the MLGs differed at multiple loci or at a single locus but for multiple repeats. The remaining four MLGs were near-identical to another, more frequently occurring, MLG. Of these, three comply with the assumptions for near-identicality by descent, as described in the methods section. These are ‘MLG-A2’, ‘MLG-P2’ and ‘MLG-P3’. The fourth near-identical MLG did not comply with the aforementioned assumptions and is therefore named ‘MLG-M’ rather than ‘MLG-I2’.

**Figure 2.**
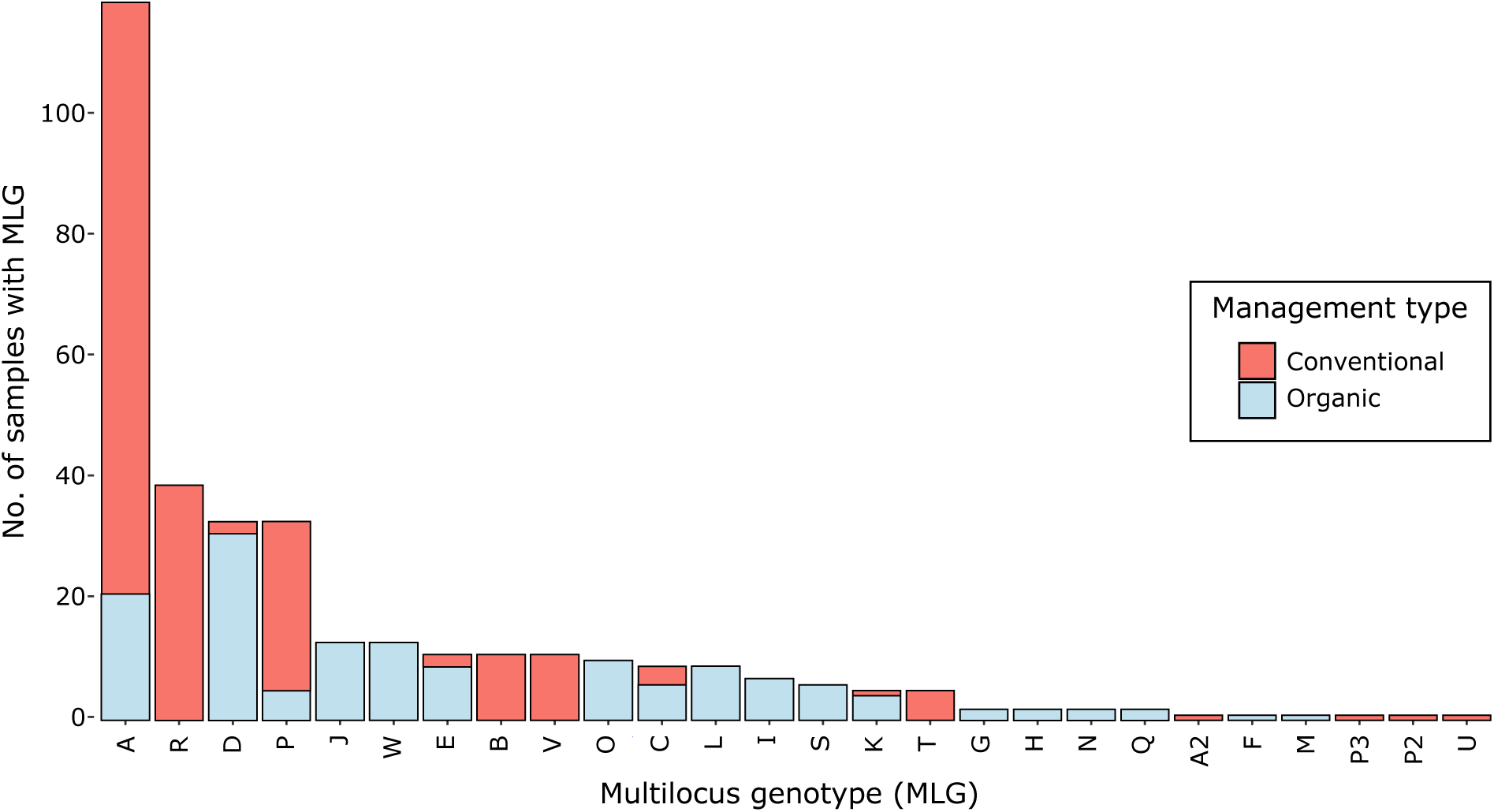
Abundance of each multilocus genotype (MLG) of *Myzus persicae*, detected in the 348 samples, collected by the researchers (large scale sampling by growers 2022 excluded) from conventional Dutch sweet pepper greenhouses in 2019-2022 and from organic greenhouses in 2019, 2020 and 2022.

Six of the MLGs were unique and detected only once, and four MLGs were detected only twice (Figure 2). The most common MLG was MLG-A, comprising 34.2% of the 348 samples. Other common MLGs were MLG-R, MLG-D and MLG-P, comprising 11.2%, 9.5% and 9.5% of the samples respectively, together making up 64.4% of all samples. Out of the total 26 MLGs, 12 were unique for organic greenhouses, eight were unique for conventional greenhouses and six were found in both greenhouse types.

Allelic richness over all samples was on average 10.4 and ranged from five for marker M37 to 15 for marker M86 (Suppl. Table S3). In organic greenhouses, allelic richness was 9.2 on average and ranged from four for marker M37 to 15 for marker M86, whilst in conventional greenhouses allelic richness was 8.4 on average and ranged from four for marker M37 to 11 for markers M86 and myz9. The average observed heterozygosity (*H_o_*) was 0.92 (ranging from 0.67 – 1) for samples from organic greenhouses, and 0.86 (0.64 – 1) for samples from conventional greenhouses.

#### Hardy-Weinberg equilibrium and linkage disequilibrium

None of the markers, for neither of the populations, was in HWE (Suppl. Table S3). Additionally, significant linkage was found among markers both for the organic (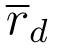 = 0.393, *p <* .001) and the conventional population (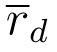 = 0.574, *p <* 0.001) without, as well as with clone-correcting the data (org: 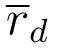 = 0.253, *p <* 0.001; conv: 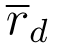 = 0.175, *p <* 0.001). In all cases, the strongest association was between markers M86 and myz9, which ranged from 0.438 for the clone-corrected organic population to 0.839 for the original conventional population (for the 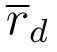 of all pairs of loci, see Suppl. Table S5).

#### MLGs in organic and conventional greenhouses in 2019 and 2020

We started by analysing the samples obtained from both organic and conventional greenhouses in 2019. We often detected multiple MLGs present at the same sampling point in a single organic greenhouse, while this rarely happened for conventional greenhouses. To better capture the richness in MLGs in the organic greenhouses, we analysed additional samples from the pool of originally sampled aphids. Consequently, more samples were analysed from organic greenhouses (N = 116) than from conventional ones (N = 63).

Great differences in population genetic structure were observed between the two different greenhouse types in 2019 as more MLGs were found in organic versus conventional greenhouses (17 vs. five respectively; Figure 3; Suppl. Table S4). On average, a significantly higher number of MLGs were simultaneously detected during a single sampling time period in an organic greenhouse (2.33 ± 0.27; average ± standard error) than in a conventional greenhouse (1.10 ± 0.07; Wilcoxon’s signed-rank test, W = 99, *p <* 0.001; Suppl. Table S6). Additionally, the specific MLGs present in an organic greenhouse changed over the different sampling periods while the same MLG was found in a conventional greenhouse over the three sampling periods (Figure 3). The average number of MLGs detected in an organic greenhouse during the three sampling periods of 2019 combined was 4.0 ± 0.58 versus 1.2 ± 0.13 for a conventional greenhouse (Wilcoxon’s signed-rank test, W = 83, *p* = .001; Suppl. Table S7).

**Figure 3.**
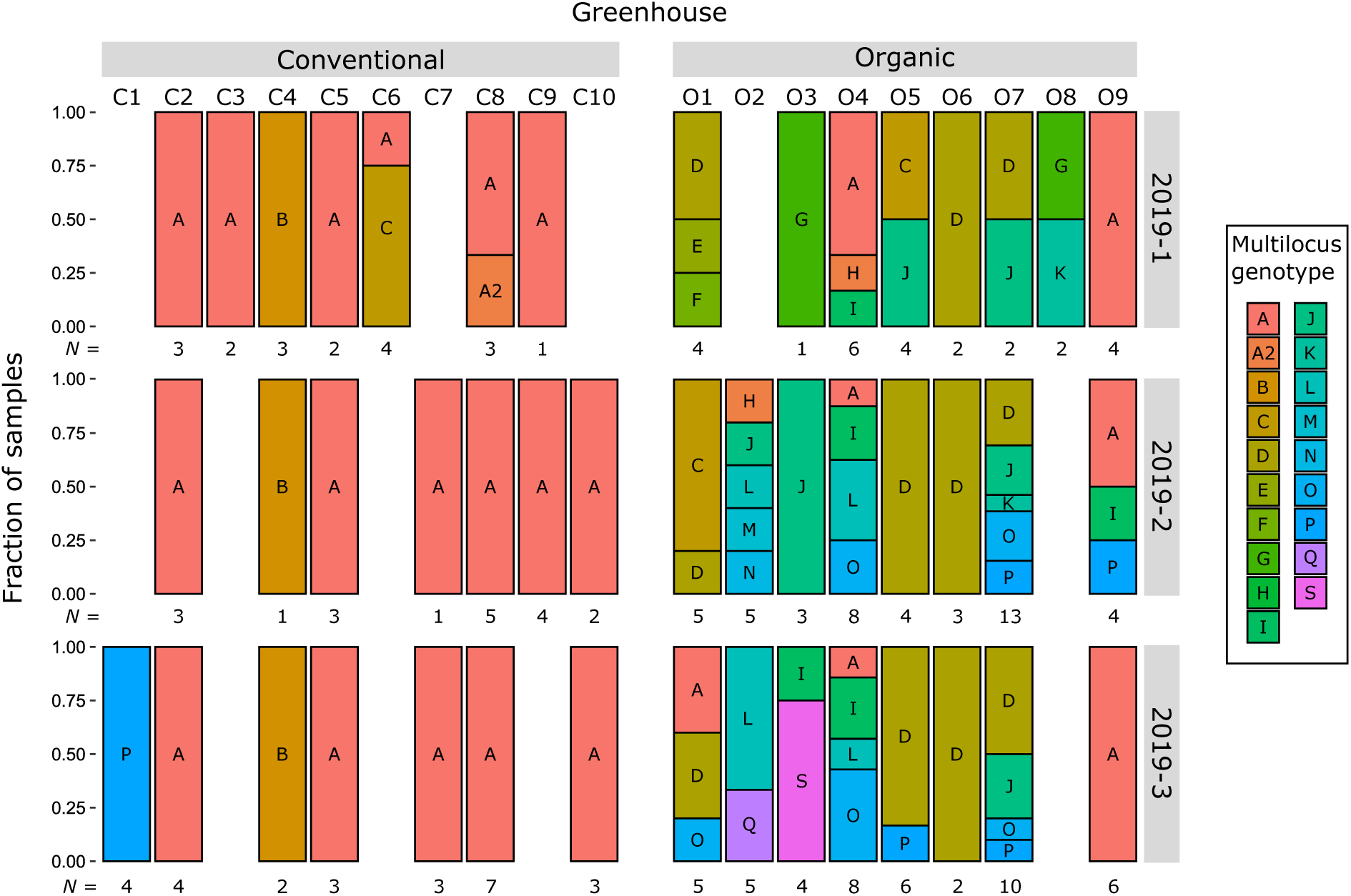
The population genetic structure of *Myzus persicae* in conventional versus organic sweet pepper greenhouses, sampled during three intermittent time periods in 2019. Every column represents the population in a greenhouse during a specific sampling time period. Every colour represents a unique multilocus genotype (MLG), determined with the use of five microsatellites. Every greenhouse identifier represents a unique greenhouse, with those starting with ‘C’ being conventional and those starting with ‘O’ being organic. Not every greenhouse could be sampled during all of the time periods due to either the crop being removed halfway during the growth season (greenhouse identifier O8) or no aphids being present (all other missing data points). The number of aphids analysed per greenhouse is displayed as ‘*N* = x’

In conventional greenhouses in 2019, MLG-A was highly dominant, comprising 49/63 (77.8%) of all genotyped samples from this greenhouse-type (Figure 3). This dominant MLG was detected in 8/10 conventional greenhouses and during all of the three sampling time periods. Other MLGs detected in this greenhouse type in 2019 were MLG-A2, MLG-B, MLG-C, and MLG-P. During only two sampling time periods we observed two different MLGs simultaneously (Figure 3). However, in one of these cases the second MLG was near-identical to the first.

In organic greenhouses in 2019, the most dominant MLG was MLG-D which was found in 31/116 (26.7%) of all genotyped samples from this greenhouse type, although it was only detected in 4 out of 9 organic greenhouses (Figure 3). The next most abundant MLGs were MLG-A, detected in 21/116 (18.1%) samples, and MLG-J, detected in 13/116 (11.2%) samples. Altogether, many different MLGs were observed in organic greenhouses in 2019, both within (ranging from one to five during a single sampling time period, and from one to six for all three sampling time periods combined) and between greenhouses (Figure 3).

At the beginning of 2020, five samples from conventional greenhouses (five greenhouses, one aphid sample each) and four samples from three organic greenhouses were sent to us by growers by post. In the organic samples we detected MLGs K, N, and Q, which were also detected in 2019, although in different greenhouses (Suppl. Table S4).

#### A change in dominant MLG in conventional greenhouses over subsequent years

*Myzus persicae* with MLG-A was highly dominant in conventional sweet pepper greenhouses in 2019 (Figure 3). Therefore, we continued sampling conventional greenhouses to determine whether MLG-A continued to dominate in this greenhouse type over time, as this could indicate positive selection for this MLG under this specific type of pest management strategy.

We observed a decline in the proportion of samples with MLG-A in conventional greenhouses over the subsequent years (Figure 4). By the end of 2021, MLG-A was detected in only a small proportion of the samples, and a new dominant MLG, MLG-R, emerged, which had not been observed before. In 2022, MLG-A was no longer detected and MLG-R became highly dominant.

**Figure 4.**
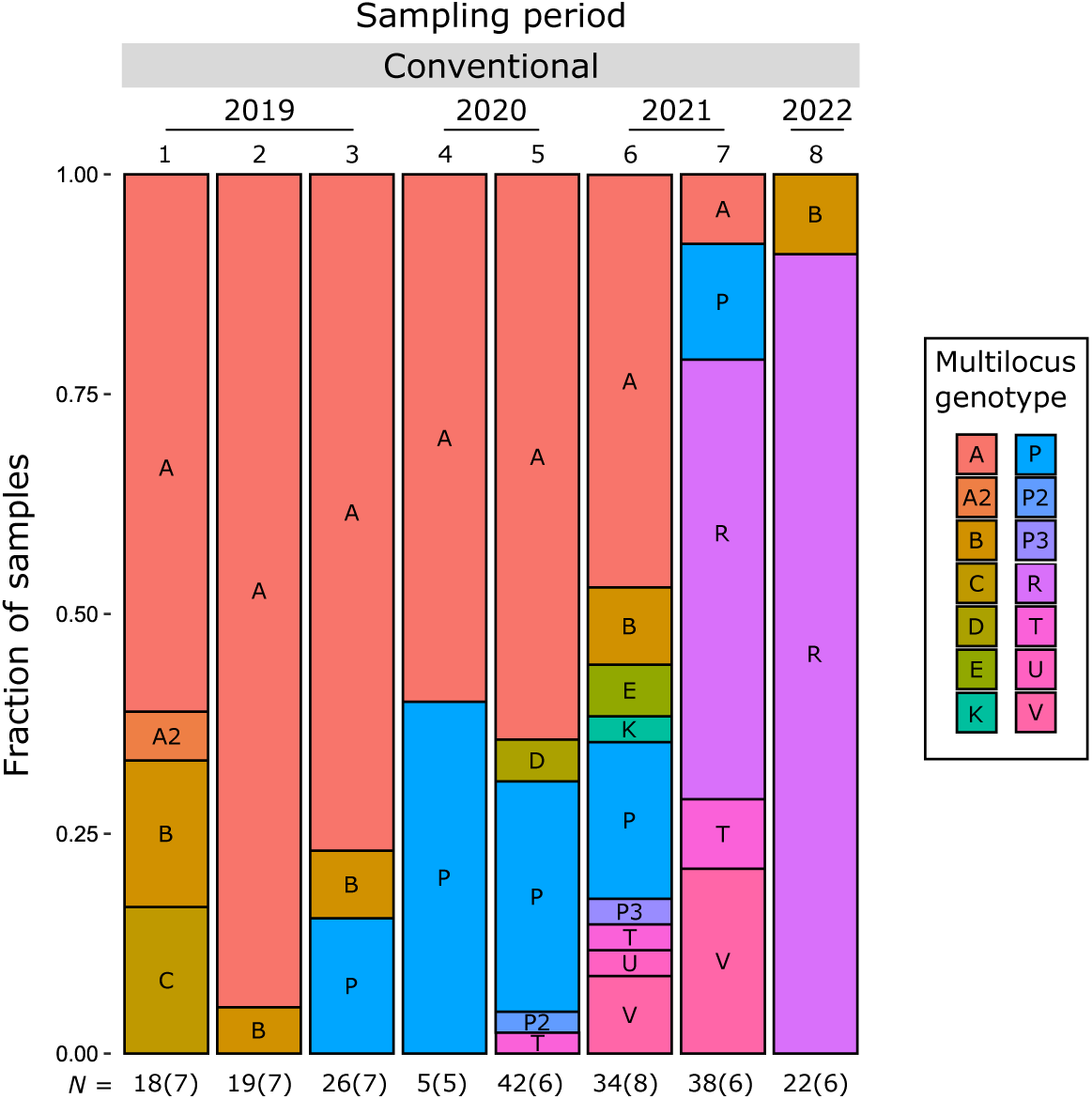
The population genetic structure of *Myzus persicae* in conventional Dutch sweet pepper greenhouses, sampled from 2019-2022, over eight sampling periods. This data clearly illustrates the decline of the dominant multilocus genotype (MLG-A) from 2019 onwards, which is subsequently replaced by a new MLG, MLG-R, by the end of 2021. The represented data includes all samples collected by the researchers, as well as samples sent in by growers in 2020, but excludes those sent in by growers in 2022. Sample sizes are displayed as ‘*N* = x(y)’ where x equals the number of aphids analysed this period and y represents the number of greenhouses sampled this period.

Of the five samples received from conventional greenhouses at the beginning of 2020, three belonged to MLG-A and the others were MLG-P (Figure 4; Suppl. Table S4). From August/September 2020 onwards, greenhouses were sampled again by the researchers and we analysed 42 samples collected from six conventional greenhouses during this time period. At that time, 27/42 (64.3%) samples were identified as MLG-A. Four greenhouses had only *M. persicae* with MLG-A and two greenhouses had MLG-A combined with multiple other MLGs. On average, during the second sampling time period in 2020, 1.83 ± 0.54 (average ± standard error) MLGs were found per conventional greenhouse. In May/June 2021 we analysed 31 samples from seven greenhouses, from which 13 (41.9%) were MLG-A, and an additional eight other MLGs were detected. Four of the greenhouses each harboured a single, distinct MLG. During this time period, on average 1.88 ± 0.52 MLGs were found per greenhouse. In September 2021, 38 samples were analysed from six greenhouses. During this time period, only 3/38 samples were MLG-A and 19/38 (42.1%) samples had a new MLG, MLG-R, which had not been detected before. This new MLG was detected in five out of the seven sampled greenhouses. Here, an average of 2.17 ± 0.40 MLGs were found per greenhouse. In 2022, 22 samples from six greenhouses were analysed: MLG-A was not detected, and 20/22 (90.1%) samples were of the new MLG-R. At this moment, 1.17 ± 0.17 MLGs were found on average per conventional greenhouse. There was a noticeable trend in the average number of MLGs detected per conventional greenhouse between the different time periods, with more MLGs in 2020 and 2021 compared to those in 2019 and 2022. However, this increase was not statistically significant (Kruskal-Wallis test, H(6) = 11.46, *p* = 0.075; Table 1).

**Table 1.** Average number of MLGs detected per greenhouse. Greenhouses were sampled during Feb/March, May-July and Sept/Oct 2019 (sampling time periods 1-3), May and Aug/Sept 2020 (sampling time periods 4-5), May/June and Sept 2021 (sampling time periods 6-7) and during March and May 2022 (sampling time periods 8-9). For example, during sampling time periods two and three, every conventional greenhouse only harboured a single MLG. These results differ from those shown in Figure 4 where the exact MLGs detected in all conventional greenhouses sampled during a sampling period can be seen together.

| Year | Pest management type | Sampling time period | No. of MLGs (average ± SE) |
| --- | --- | --- | --- |
| 2019 | Conventional | 1 | $1.29 \pm 0.18$ |
| 2019 | Conventional | 2 | $1.00 \pm 0.00$ |
| 2019 | Conventional | 3 | $1.00 \pm 0.00$ |
| 2020 | Conventional | 5 | $1.83 \pm 0.54$ |
| 2021 | Conventional | 6 | $1.88 \pm 0.52$ |
| 2021 | Conventional | 7 | $2.17 \pm 0.40$ |
| 2022 | Conventional | 8 | $1.17 \pm 0.17$ |
| 2019 | Organic | 1 | $1.88 \pm 0.30$ |
| 2019 | Organic | 2 | $2.75 \pm 0.62$ |
| 2019 | Organic | 3 | $2.38 \pm 0.42$ |
| 2022 | Organic | 9 | $1.80 \pm 0.37$ |

#### Large-scale analysis of the MLG-R prevalence in sweet pepper in the Netherlands

To monitor the distribution of genotypes across the Netherlands, we analysed 194 aphid samples (up to six per greenhouse) that were sampled and sent in by growers from 34 conventional sweet pepper greenhouses. MLG-R was found in 33/34 greenhouses and in 185/194 (95.4%) samples. Additionally, MLG-A was found in three samples from one greenhouse and another grower had sent six samples with MLG-D (Figure 5; Suppl. Table S2).

**Figure 5.**
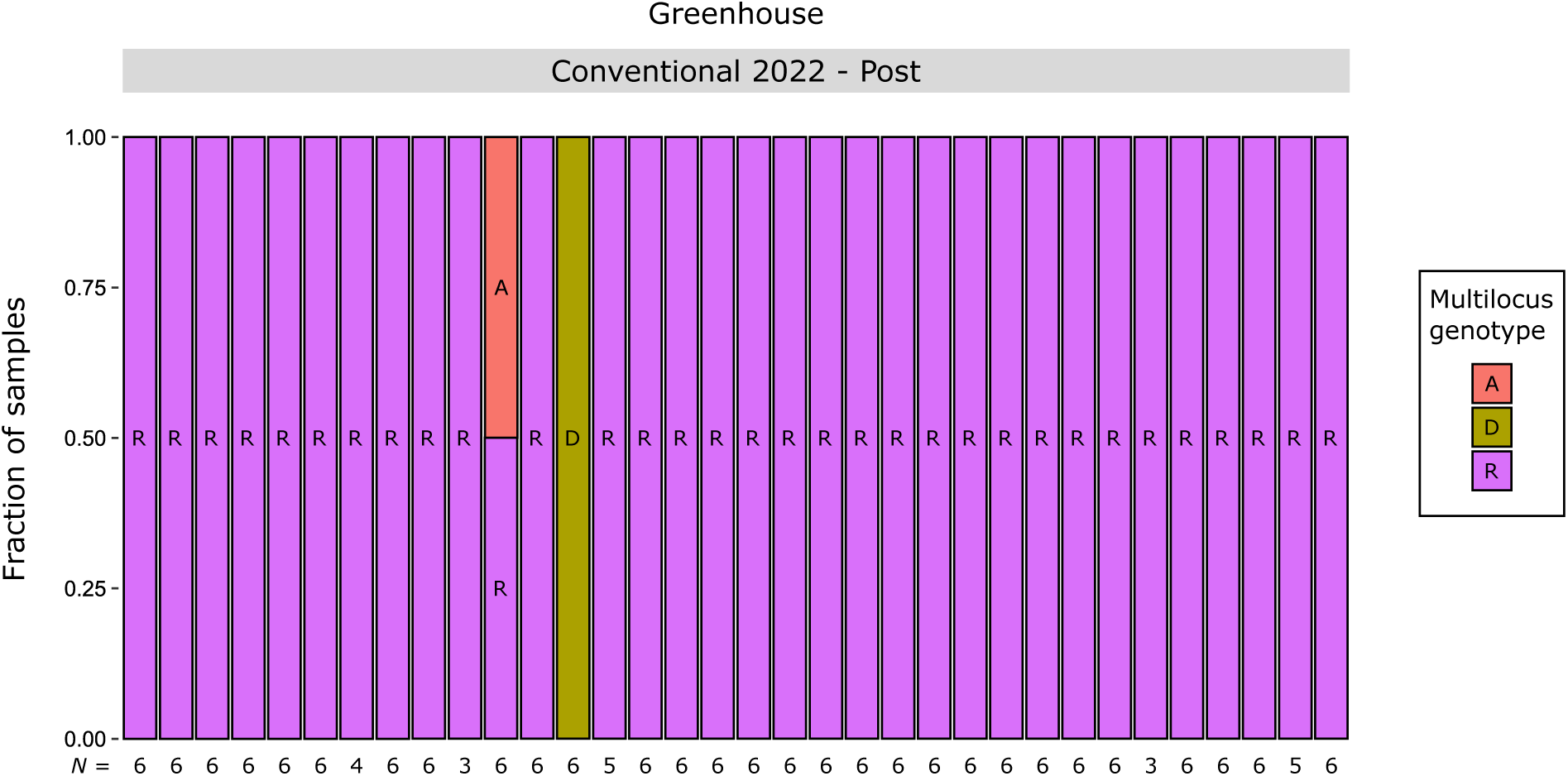
Multilocus genotypes of *Myzus persicae* collected by large scale sampling of 34 conventional Dutch sweet pepper greenhouses by growers in April 2022. Each bar represents a unique greenhouse. The number of aphids analysed per greenhouse is displayed as ‘*N* = x’

#### Questionnaire about perceived difficulties with aphid infestations and applied insecticides

The growers that sent in aphids by post in April 2022 also filled out a questionnaire. Of the 33 growers, 22 (66.7%) self-reported ‘much difficulty’ to control the aphid populations in their greenhouse at that time. Furthermore, 10 out of 33 (30.3%) self-reported ‘average difficulty’, and only one out of 33 (3.0%) ‘little difficulty’ with aphid control (Figure 6A; Suppl. Table S2). When asked about the end of their previous growth season (2021), ten out of 31 (32.3%) self-reported to have had ‘extra difficulty’ controlling aphids in their greenhouse at that time.

**Figure 6.**
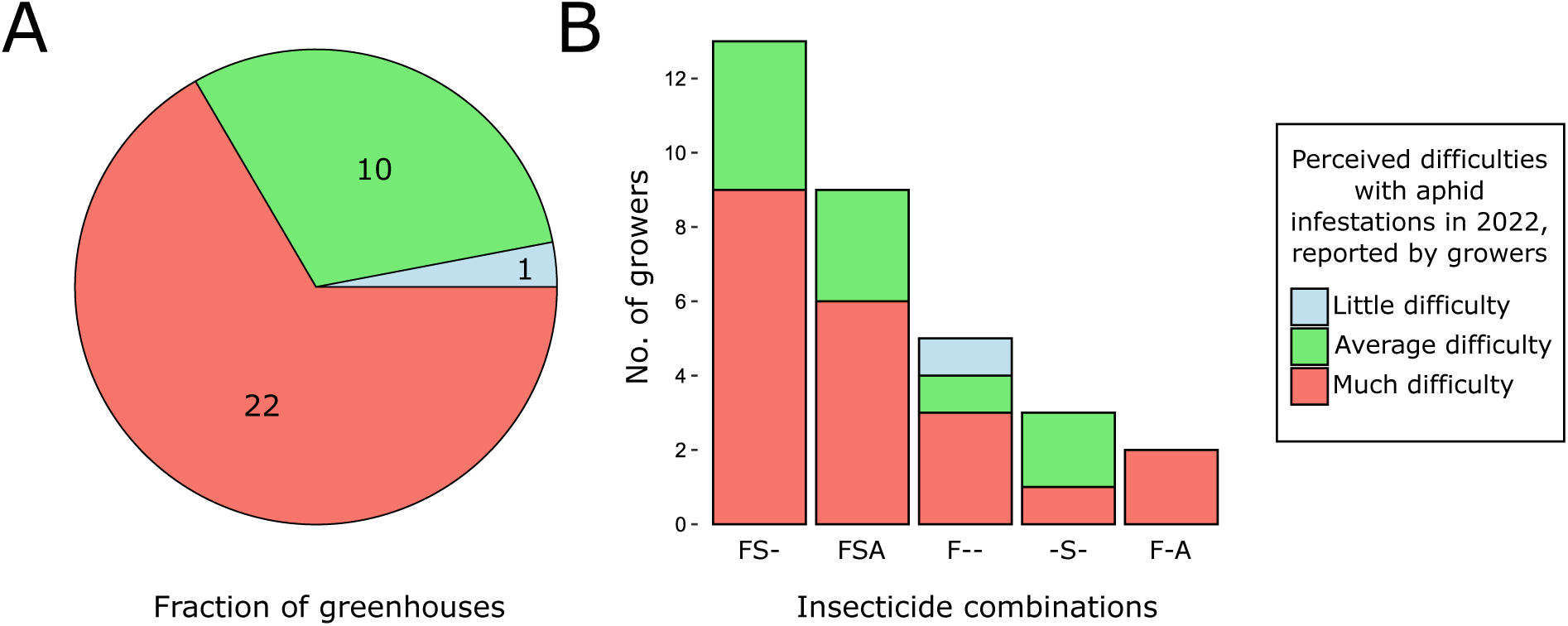
Results of questionnaire to conventional sweet pepper growers in April 2022. A) Fraction of growers self-reported to have had ‘little difficulty’ (light-blue), ‘average difficulty’ (green), or ‘much difficulty’ (red) with aphid infestations in their crop by April 2022. Numbers represent the number of growers. B) Combinations of the most commonly applied insecticides during the 2022 growth season until April 2022 where ‘F’ = flonicamid, ‘S’ = sulfoxaflor, and ‘A’ = acetamiprid.

Based on the questionnaire, the most commonly used insecticide used in the 2022 sweet pepper growth season up until April was flonicamid, which was applied in 29/32 (90.6%) of the greenhouses (Figure 6B; Suppl. Table S2). Other commonly used insecticides were sulfoxaflor, which was applied in 25/32 (78.1%) of the greenhouses, and acetamiprid, applied in 11/32 (34.4%) of the greenhouses. Most greenhouses reported to have applied both flonicamid and sulfoxaflor (13/32), or a combination of all three insecticides (9/32).

#### No MLG-R in organic greenhouses in 2022

To find out if the highly dominant MLG-R was also present in organic greenhouses in 2022, we additionally sampled five organic greenhouses (N = 24) in May 2022. In each greenhouse, one to three different MLGs (MLG-E, MLG-S, and/or MLG-Y) were detected, but never MLG-R (Figure 7; Suppl. Table S4). The MLGs detected in organic greenhouses were not detected in any of the conventional greenhouses sampled in 2022. However, the average number of MLGs detected in conventional greenhouses (1.17 ± 0.41) was not significantly different (Wilcoxon’s signed-rank test, W = 9, *p* = 0.164) from organic greenhouses (1.8 ± 0.84) in 2022 (Suppl. Table S6).

**Figure 7.**
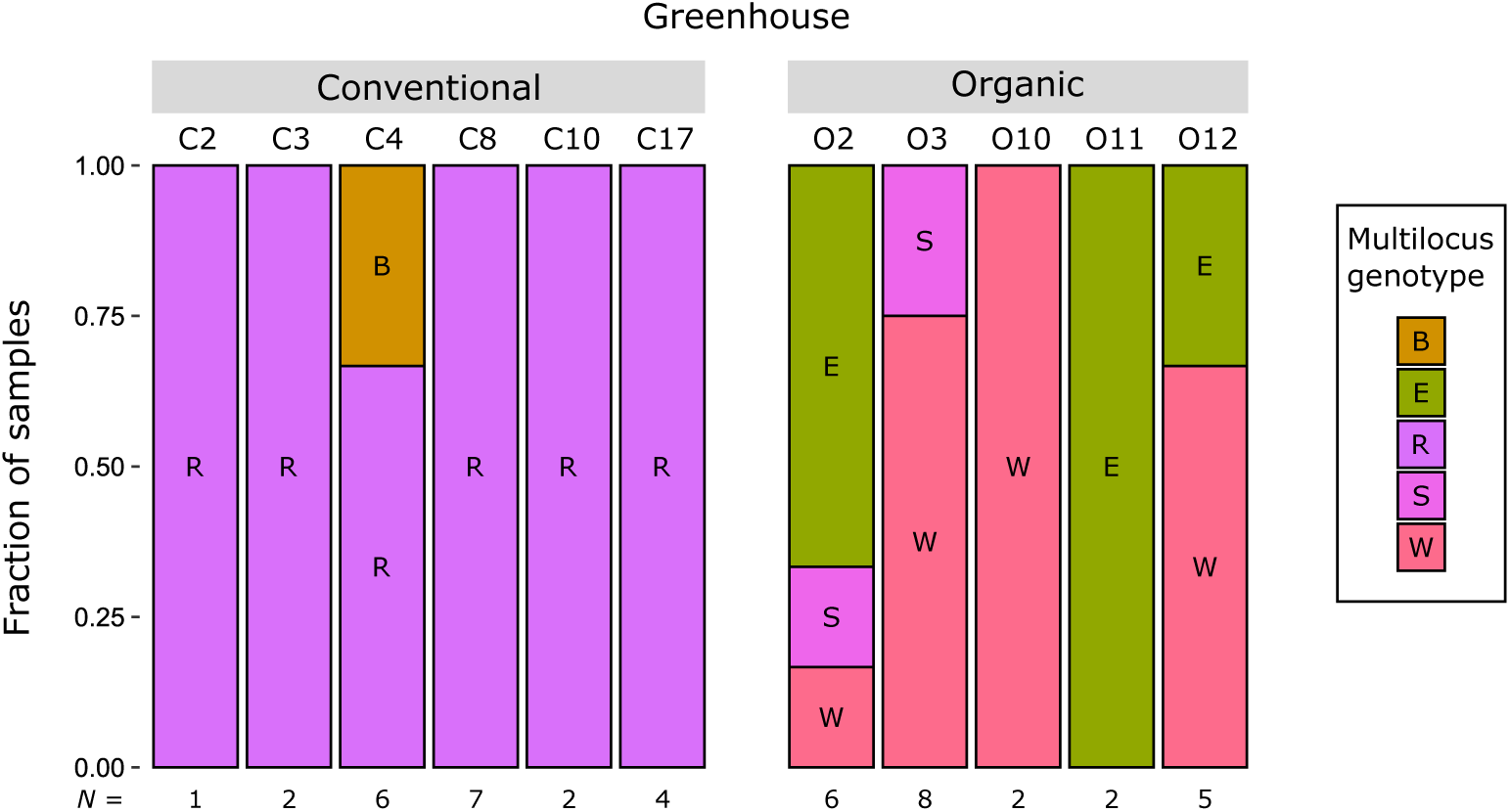
The detected multilocus genotypes (MLGs) of *Myzus persicae* in Dutch conventional and organic sweet pepper greenhouses in 2022, sampled by the researchers. Every colour represents a unique MLG, determined with the use of five microsatellites. Every greenhouse identifier represents a unique greenhouse, with those starting with ‘C’ being conventional and those starting with ‘O’ being organic.The number of aphids analysed per greenhouse is displayed as ‘*N* = x’

### Do all samples with MLG-A represent true clone-mates?

Among the 15 whole-genome sequenced aphid lines, of which seven with MLG-A and eight with other unique MLGs, 1,073,049 SNPs remained after filtering, with an average coverage of 36.9x ± 5.2 (average ± standard deviation). The overall nucleotide diversity and Waterson’s estimator (Suppl. Figure S1) averaged 1.2x10*^−^*^3^ (± 3.6x10*^−^*^4^) and 1.3x10*^−^*^3^ (± 3.7x10*^−^*^4^) respectively. Each aphid line showed higher heterozygosity levels than expected (Table 2) and on average 330,356 (± 17,472) loci were heterozygous per aphid line, which was 13,5% higher than the expected 291,011.

**Table 2.** Aphid lines for which the whole genome has been sequenced. Their ID, multilocus genotype (MLG), date and greenhouse identifier of where the aphid line was originally collected, and the sequencing run (Run) in which the *Myzus persicae* lines were whole genome sequenced. The observed number of heterozygous sites (Nhet), the inbreeding coefficient (F), and number of unique heterozygous (U(Het)) sites per aphid line. The total number of analysed sites was 1,071,088 for each aphid line.

| Aphid ID | MLG | Collection date | Greenhouse | Run | Nhet | F | U(Het) |
| --- | --- | --- | --- | --- | --- | --- | --- |
| A(C2-2019) | A | 14-Jun-19 | GH-C2 | 1 | 324654 | -0.11561 | 135 |
| A(C5-2019) | A | 19-Jun-19 | GH-C5 | 1 | 324703 | -0.11577 | 32 |
| A(O1-2019) | A | 02-Jul-19 | GH-O1 | 1 | 324854 | -0.11629 | 36 |
| A(O4-2019) | A | 19-Jul-19 | GH-O4 | 1 | 324786 | -0.11606 | 74 |
| A(C9-2019) | A | 19-Jul-19 | GH-C9 | 1 | 324540 | -0.11521 | 160 |
| A(C9-2021) | A | 28-May-21 | GH-C9 | 2 | 326364 | -0.12148 | 107 |
| A(C7-2021) | A | 07-Jul-21 | GH-C7 | 2 | 327762 | -0.12629 | 67 |
| J(O7-2019) | J | 13-Jun-19 | GH-O7 | 1 | 344375 | -0.18337 | 18,867 |
| P(O9-2019) | P | 18-Jun-19 | GH-O9 | 1 | 345424 | -0.18698 | 34,409 |
| D(O5-2019) | D | 10-Jul-19 | GH-O5 or O6 | 1 | 358626 | -0.23234 | 20,035 |
| X(O3-2019) | X | 17-Jul-19 | GH-O3 | 1 | 360257 | -0.23795 | 24,299 |
| K(C2-2021) | K | 14-May-21 | GH-C2 | 2 | 347877 | -0.19541 | 12,696 |
| T(C12-2021) | T | 28-May-21 | GH-C12 | 2 | 302125 | -0.03819 | 35,951 |
| B(C4-2021) | B | 09-Jun-21 | GH-C4 | 2 | 310014 | -0.06530 | 52,446 |
| R(C13-2021) | R | 01-Oct-21 | GH-C13 | 2 | 308980 | -0.06175 | 61,229 |

The genetic distance between aphid lines was calculated by comparing each locus in the filtered VCF file, assigning values of 0 to reference homozygotes, 1 to alternative homozygotes, and 0.5 to heterozygotes, and summing these values across all loci. The average genetic distance between aphid lines with different MLGs was 266,663 (± 35,899), which is significantly higher than the average of 2,920 (± 879) between aphid lines with MLG-A. Hierarchical clustering revealed that all aphid lines with MLG-A clustered closely together compared to all other aphid lines (Figure 8). The number of unique mutations in het-erozygous state ranged from 32 to 160 for MLG-A aphid lines, while for all other aphid lines, it varied between 12,696 and 61,229 sites. Notably, 51,430 unique heterozygous sites were shared between the various MLG-A lines, but were fixed in one homozygous state in all other aphid lines. This high number of shared heterozygous sites but low number of unique variants among MLG-A aphid lines, compared to any pairwise comparison between aphid lines of other MLGs (Figure 9), suggests that they are true clone-mates, all originating from a single parthenogenetic female that resulted from a sexual cross.

**Figure 8.**
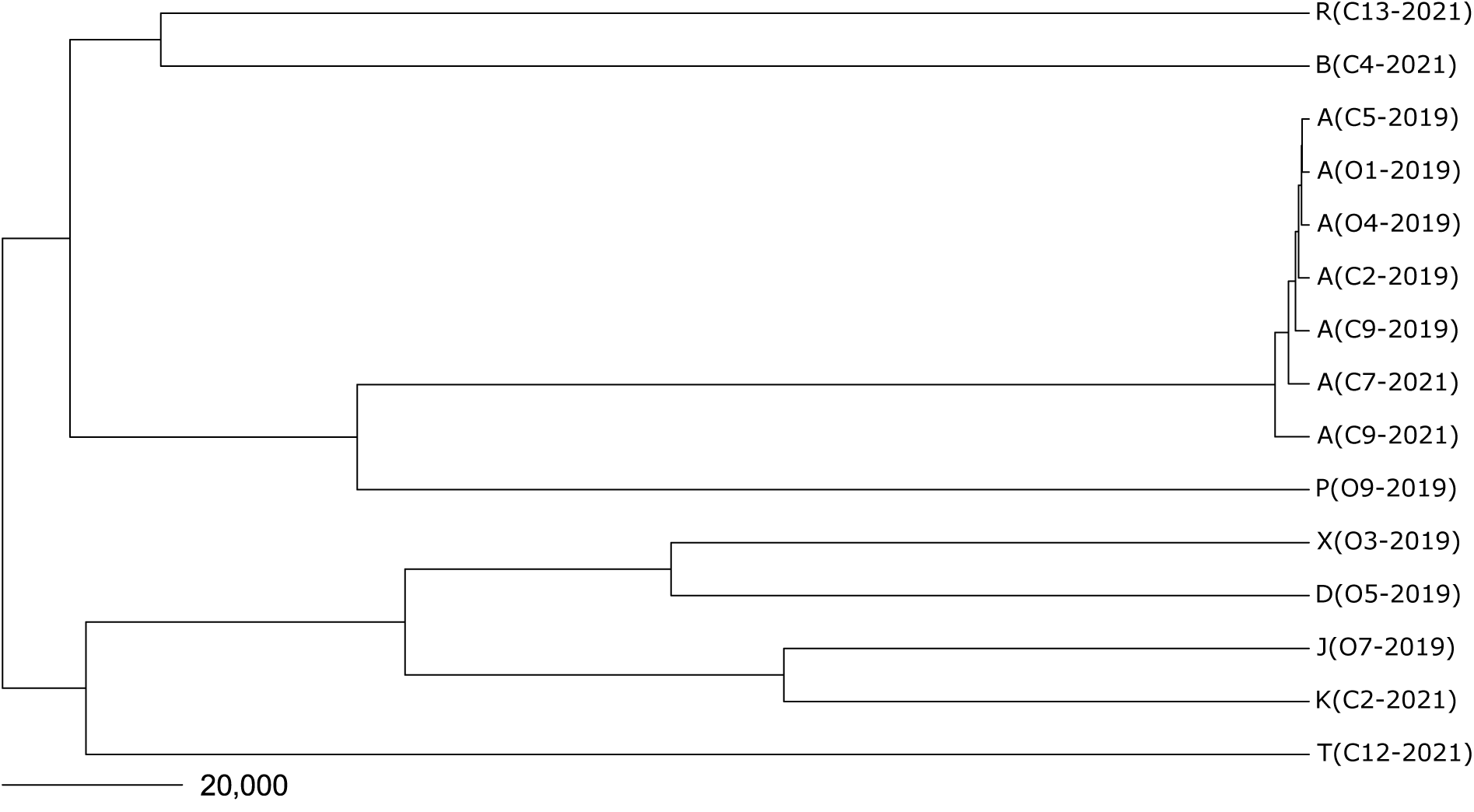
Hierarchical clustering of the aphid lines for which the whole genome has been sequenced, based on genetic distance, whereby loci that show different homozygous states are coded as a distance of 1, and a difference between homozygous and heterozygous state as 0.5. The scalebar shows the length of a branch that indicates a genetic distance of 20,000.

**Figure 9.**
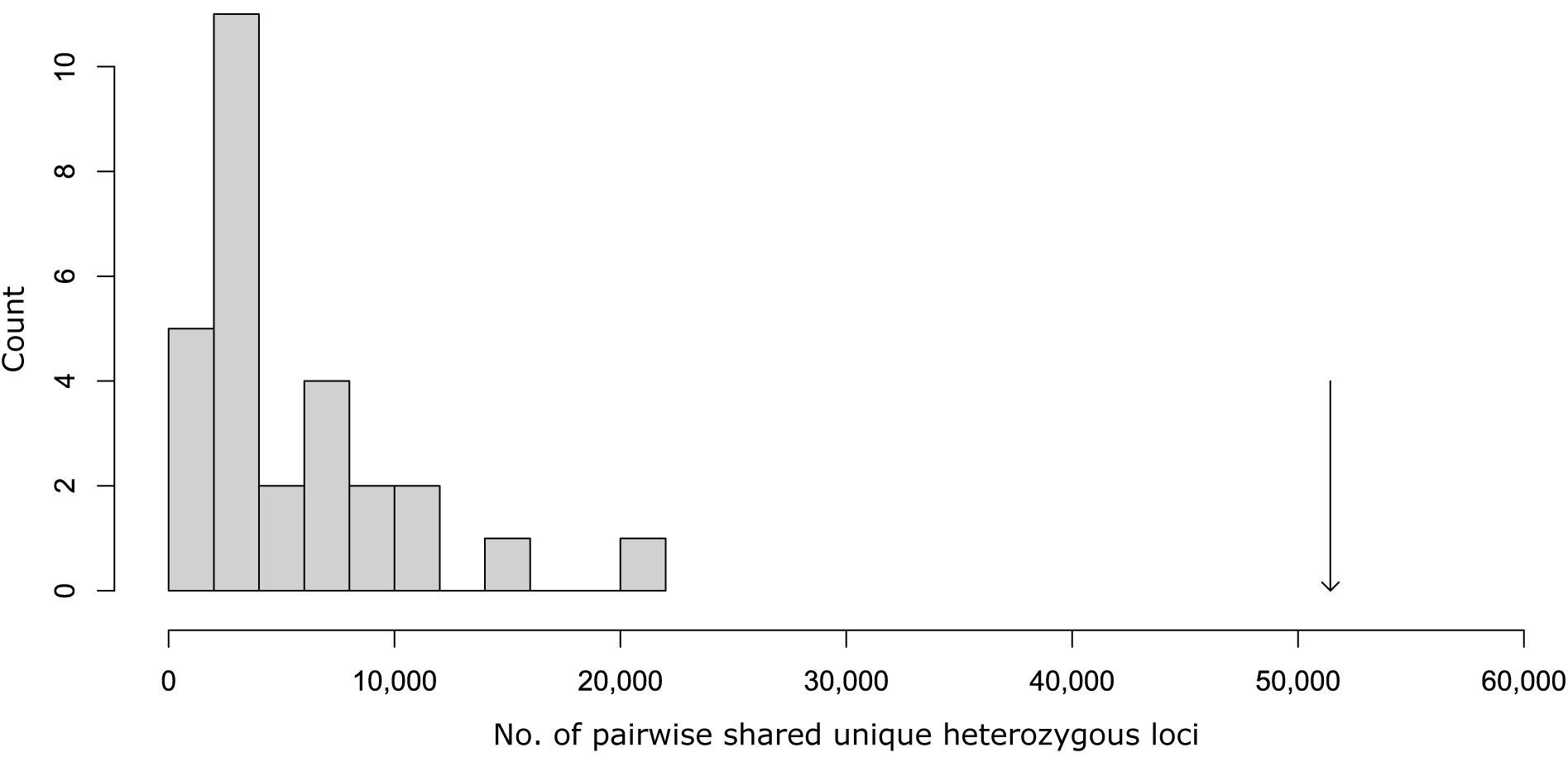
Distribution of number of pairwise shared unique heterozygous loci between all whole genome sequenced aphid lines except those with MLG-A. The number of unique heterozygous loci shared among all MLG-A lines is depicted by the arrow

Furthermore, other pairs of aphid lines that were more closely related, based on the hierarchical clustering, also exhibited a high number of shared heterozygous sites. This was for instance the case for J(O7-2019) and K(C2-2021) with 20,481 sites, D(O5-2019) and Z(O3-2019) with 10,665 sites, and R(C13-2021) and B(C4-2021) with 15,102 sites. These findings support the validity of the hierarchical clustering method, as it successfully identified more closely related aphid lines, which have more shared heterozygous sites due to their more recent common ancestors.

## Discussion

The strong selection pressures exerted by pest control strategies in commercial greenhouses can lead to the rapid evolution of resistant pest populations, both due to conventional and organic pest control methods (Mota-Sanchez and Wise, 2023; Siegwart et al., 2015; Tabashnik and Carrière, 2015; Tomasetto et al., 2017). Successful control of *Myzus persicae*, the economically most problematic aphid species worldwide, in sweet pepper greenhouses remains inconsistent (Glastuinbouw Nederland, 2022; Messelink et al., 2011). Because *M. persicae* reproduces clonally through apomictic parthenogenesis on its secondary hosts, clonal amplification is expected for genotypes that are especially successful in the commercial greenhouse environment. In this study, we explored the clonal diversity and population genetic structure of *M. persicae* in organic and conventional Dutch sweet pepper greenhouses over multiple consecutive years with the use of five microsatellites. Accordingly, we investigated whether specific clonal lines dominate, as this could indicate adaptation of the pest populations to pest control methods. We observed very clear differences in population genetic structure, as specific MLGs clearly dominated in conventional greenhouses, but no dominating MLGs were observed in organic greenhouses. These results suggest that conventional pest control selects more strongly for specific aphid genotypes than organic pest control.

### Population genetics of *Myzus persicae* in Dutch sweet pepper

Microsatellites are useful genetic markers to study the genetic structure of aphid populations due to their relatively high mutation rate and the high level of polymorphism of individual loci. Consequently, a few loci are already enough to distinguish distinct clonal lines. Here, we used five previously developed microsatellite markers (Sloane et al., 2001; Wilson et al., 2004) and detected a total of 26 MLGs in 348 samples. Strikingly, only four MLGs comprised approximately two thirds of the samples. This low clonal diversity and the dominance of specific (time-persistent) clones in populations of *M. persicae* have been described before and is most often attributed to either the presence of the population on secondary hosts on which only parthenogenetic reproduction takes place (Sanchez et al., 2013), the absence of the primary host resulting in an anholocyclic life cycle (Fenton et al., 2005), and/or high proportions of obligate parthenogens (Guillemaud et al., 2003; Sanchez et al., 2013; Vorburger et al., 2003; Zamoum et al., 2005), which is linked to climatic conditions (Rispe et al., 1998; Rispe and Pierre, 1998). Furthermore, the presence of insecticide-resistant clones under insecticide treatment has been shown to lead to low clonal diversity and the dominance of specific clones (Fenton et al., 2005; Roy et al., 2013). However, one should consider that clonal diversity will also depend on the rate of influx of new genotypes into a system, which can be expected to be lower in semi-closed systems, such as the greenhouses we sampled in this study, compared to open agricultural fields and natural areas where most previous studies have sampled.

Besides low clonal diversity and the dominance of specific clones, we also observed severe deviations from HWE, significant linkage among loci, and high levels of heterozygosity, which all indicate an anholocyclic life cycle (Balloux et al., 2003; Halkett et al., 2005). This is to be expected for *M. persicae* in the absence of its primary host, or in climatically homogeneous environments as is the case in sweet pepper greenhouses. Notably, higher levels of heterozygosity in a population have been previously linked to higher proportions of obligate parthenogenetic clones in *M. persicae* (Vorburger et al., 2003). Thus, potentially (part of) the *M. persicae* population in Dutch sweet pepper greenhouses consists of obligate parthenogenetic lineages. However, additional experiments are needed to reveal the exact reproductive mode of each sampled line of *M. persicae*.

### Whole genome sequencing

We used five microsatellite markers, which is fewer than most former studies genotyping *M. persicae*, using six (Hlaoui et al., 2019; Roy et al., 2013), seven (Guillemaud et al., 2003; Vorburger et al., 2003; Zepeda-Paulo et al., 2010), or eight (Sanchez et al., 2013; Zamoum et al., 2005) markers. Although Fenton et al. (2005) used four microsatellite markers, they augmented these with intergenic spacer fingerprints to obtain additional genetic information. To verify that the persistent observation of MLG-A was genuinely due to a high prevalence of a single clonal line, we conducted whole genome sequencing (WGS) on 15 aphid lines, of which seven with MLG-A.

The genetic diversity observed in our *M. persicae* lines is comparable to that of *Nasonia vitripennis* (Pannebakker et al., 2020) and within one order of magnitude of *Drosophila melanogaster* wild populations (Kapun et al., 2020). Additionally, our WGS results revealed an excess of heterozygosity, consistent with the microsatellite results. Notably, the genetic distance between the MLG-A lines collected during different years and from various greenhouses was small, with an average of 2,920 (± 879), compared to 266,663 (± 35,899) between lines with different MLGs. The substantial number of shared unique heterozygous sites between the various MLG-A lines (51,430; Figure 9) suggests that all MLG-A samples originated from a single parthenogenetic female that resulted from a sexual cross. Therefore, we conclude that five microsatellite markers were sufficient to distinguish the unique clonal lines in this study and a good starting point for analysis in general. However, we acknowledge that additional markers may be necessary when studying populations under conditions of sexual reproduction, or in open systems where the influx of new genotypes is higher.

In 1977, Janzen proposed that aphid clonal lines are actually super-organisms, with all individual clone-mates representing one “evolutionary individual” (Janzen, 1977). This view was later rejected as it became clear that mutational events induce genetic variation between clone-mates (Loxdale, 2008). Our WGS results confirm this, showing that despite all *M. persicae* lines with MLG-A originated from a single parthenogenetic female that resulted from a sexual cross, none of the lines were 100% genetically identical.

### Population genetic structure in conventional greenhouses

In conventional greenhouses, we initially observed the dominance of a single MLG, MLG-A, which was later almost entirely replaced by the ‘new’ MLG-R, after a short phase of higher clonal diversity. The high prevalence of these two clones could hypothetically be attributed to generally higher reproductive rates compared to other clones (Herzog et al., 2007; Vorburger, 2005). However, the low prevalence of MLG-A in organic greenhouses throughout the years, and the absence of MLG-R in organic greenhouses in 2022, rather suggest the involvement of pest management strategy-specific factors.

One important distinction in applied pest control methods between organic and conventional greenhouses lies in the use of chemical insecticides, which are exclusively used in conventional crop systems. Before its EU ban in 2020, the primary insecticide for controlling *M. persicae* in Dutch sweet pepper was pymetrozine (Beekman et al., 2022; Ctgb, 2024). After the ban on pymetrozine, growers transitioned to using flonicamid, which was approved for use against *M. persicae* in Dutch sweet pepper since 2018 (Ctgb, 2024). The switch from pymetrozine to flonicamid in 2020 coincides with reduced prevalence of MLG-A in conventional greenhouses from 2020 onwards. This is accompanied by increased clonal diversity and may indicate that MLG-A has a reduced sensitivity to pymetrozine. Likewise, the sudden emergence and subsequent success of MLG-R in 2021 and 2022 could indicate strong positive selection for this new genotype under the new selective environment. Thus, the success of MLG-R may be linked to reduced sensitivity to flonicamid. This could explain the reported challenges in controlling *M. persicae* by growers in 2022.

Having never been detected before, MLG-R was found for the first time during the second sampling time period of 2021, which occurred in the third trimester of the crop growth season. At that time, MLG-R was detected in five out of seven sampled conventional greenhouses. This sudden emergence raises the question of how new genotypes are introduced into greenhouses. One possibility is that genotypes are hitchhiking on new plants distributed by plant propagators, potentially introducing a founder effect. Alternatively, genotypes may originate from natural populations or other crop systems and enter the greenhouse as winged individuals, known as alata, which are produced in response to crowding (Shaw, 1970; Sutherland, 1969). In 2019, every conventional greenhouse we sampled sourced their plants from a different propagator (unpublished data), and we assume this also happened during the subsequent years. Furthermore, MLG-R was not detected in any of the sampled conventional sweet pepper greenhouses during the first sampling time period of 2021. Therefore, we can reasonably rule out that this genotypes was initially distributed by a plant propagator. We hypothesise that MLG-R entered greenhouses in 2021 at some time during the crop growth season as alata, after which positive selection for this MLG, potentially driven by insecticide application, could have increased their frequency in conventional greenhouses. After initial settlement in a crop system, further spread of an MLG by plant propagators in the subsequent years is plausible.

To validate the above-stated hypotheses, future research should assess MLG-A and MLG-R’s sensitivity to pymetrozine, flonicamid, and potentially also to other insecticides used in sweet pepper, comparing them to less prevalent genotypes from sweet pepper.

### Population genetic structure in organic greenhouses

Contrary to what we observed in conventional greenhouses, no specific MLGs dominated in organic sweet pepper greenhouses in 2019. Biocontrol methods applied in these organic greenhouses involve the deployment of parasitoid wasps, aphid predators such as *Aphidoletes aphidimyza*, and the use of plant-based bio-insecticides such as Neem-based products (Beekman et al., 2022). Resistance to parasitoids is hypothesized to potentially hamper aphid control in organic greenhouses (Vorburger, 2018), as previous laboratory studies have shown that aphids carrying heritable protective secondary endosymbionts may outcompete aphid lineages without such endosymbionts under parasitoid pressure (Herzog et al., 2007; Käch et al., 2018; Oliver et al., 2008). Additionally, aphids display varying levels of endogenous resistance to parasitoids (Martinez et al., 2014; Sandrock et al., 2010; Von Burg et al., 2008). However, the 2019 samples genotyped in this study were shown be free of secondary endosymbionts (Beekman et al., 2022), and no differences in sensitivity to the biocontrol parasitoids *A. matricariae* and *A. colemani* could be observed between nine *M. persicae* lines collected from the Dutch sweet pepper greenhouses in 2019 (Suppl. Table S8 displays how those aphid lines are named in this study). The absence of dominating genotypes in organic greenhouses in 2019 means that no specific genotypes outcompeted others under biocontrol practices, which suggests that none of the detected genotypes is resistant to the combined biocontrol practices applied in these systems.

The results in 2022 were different from those of 2019, and limited genotypic variation was observed in the five sampled organic greenhouses, with only three MLGs (E, S, and W). It is plausible that these, particularly MLG-E and MLG-S, which were previously detected in 2019 but in low frequencies, increased in frequency over time due to positive selection or genetic drift. However, the absence of data from organic greenhouses in 2020 and 2021 precludes making definitive conclusions. A more extensive study over space and time would be necessary to confirm potential positive selection for MLGs E, S and/or W under organic pest management.

### Potential consequences of population genetic structure differences between organic and conventional systems

Although we found no proof for positive selection on certain *M. persicae* genotypes under organic pest management, the presence of higher genetic diversity in populations of *M. persicae* in organic greenhouses can be expected to influence the efficacy of biocontrol agents differently in organic versus conventional systems. For instance, our results imply that biocontrol agents deployed in organic greenhouses will need a wider range of traits to handle potential resistance mechanisms encoded by the various aphid holobionts (host plus associated microbes; Bordenstein and Theis, 2015) in these systems. Adapting to diverse host genotypes may involve a switching penalty, such as increased searching and handling time. Previous work showed that a switching penalty in parasitoid wasps can enable the coexistence of various aphid genotypes, such as those with and without protective endosymbionts (Preedy et al., 2020). Even though previous work showed no facultative endosymbionts and no variation in endogenous resistance to parasitoids in *M. persicae* from Dutch sweet pepper greenhouses (Beekman et al., 2022), variation in sensitivity and/or protective behaviours towards other biocontrol agents are still possible. In conventional systems, on the other hand, biocontrol agents would ideally be optimised for their effectiveness towards the dominant genotypes. However, since the dominant aphid genotype can change over time, as we have demonstrated, it will be necessary to closely monitor which genotype is currently dominant.

## Conclusion

With this study, we are the first to demonstrate the impact of different pest management strategies (conventional versus organic) on the population genetic structure of *M. persicae* within commercial greenhouses. Our findings reveal that monitoring the population genetic structure of this important crop pest can provide valuable insights that may help predict outbreaks and can possibly even signal the emergence of resistance to the applied pest control methods. While genetic monitoring of pest populations for the purpose of improving pest control success is currently underutilised, our research underscores its potential in developing and optimising sustainable and efficient pest management strategies in agricultural settings.

## Acknowledgements

We are very grateful to Bert Dibbits and Kimberley Laport for performing the fragment analyses. We are grateful to all participating growers for letting us sample in their greenhouses, and for filling out the questionnaire and sending aphid samples by post. We would like to thank Christoph Vorburger for his advice on microsatellite analyses. We are grateful to Tom Groot for his role in enabling the collaboration with Glastuinbouw Nederland. We would like to thank Jeannette Vriend and Joke Vreugdenhil for their involvement in the large-scale sampling effort. Lastly, we would like to thank Christoph Vorburger, Tom Groot, Tibor Bukovinszky, Ben Philip, and Gerben Messelink for the insightful discussions.

## Data availability statement

Supplementary tables and supplementary file S1 are available on figshare and are accessible via the following link: https://doi.org/10.6084/m9.figshare.26145952. Raw sequence data have been deposited in the Sequence Read Archive (http://www.ncbi.nlm.nih.gov/sra) under the BioProject number PRJNA1065815.

## CRediT authorship contribution statements

**Mariska M. Beekman:** Conceptualisation, Formal analysis, Funding acquisition, Investigation, Methodology, Visualisation, Writing – original draft. **Joost van den Heuvel:** Formal analysis, Visualisation, Writing - Original Draft. **S. Helena Donner:** Investigation, Writing - Review & Editing. **Jordy J. H. Litjens:** Resources, Investigation. **Marcel Dicke:** Conceptualisation, Supervision, Writing - Review & Editing. **Bas J. Zwaan:** Conceptualisation, Supervision, Funding acquisition, Writing - Review & Editing. **Eveline C. Verhulst:** Conceptualisation, Funding acquisition, Methodology, Supervision, Writing - Review & Editing. **Bart A. Pannebakker:** Conceptualisation, Funding acquisition, Methodology, Project administration, Supervision, Writing - Review & Editing.

## Supplementary Files

### Supplementary Tables

All supplementary tables are accessible via the following link: https://doi.org/10.6084/m9. figshare.26145952

Table S1: Information on all greenhouses where aphids were sampled between 2019 and 2022, including sample sizes.

Table S2: Information on the greenhouses whose growers voluntarily participated in a large-scale sampling effort in April 2022.

Table S3: Information on the microsatellite markers, including their characteristics, testing statistics, primer sequences, and sources.

Table S4: Results of multilocus genotypes of *Myzus persicae* samples, sampled by researchers from 2019-2022

Table S5: The 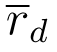-values between all pairs of microsatellite loci for raw and clone-corrected data.

Table S6: The number of different MLGs detected in each greenhouse per sampling period.

Table S7: The total number of different MLGs that were found in a greenhouse in 2019.

Table S8: Aphid lines: Identifiers and additional information.

**Figure S1.**
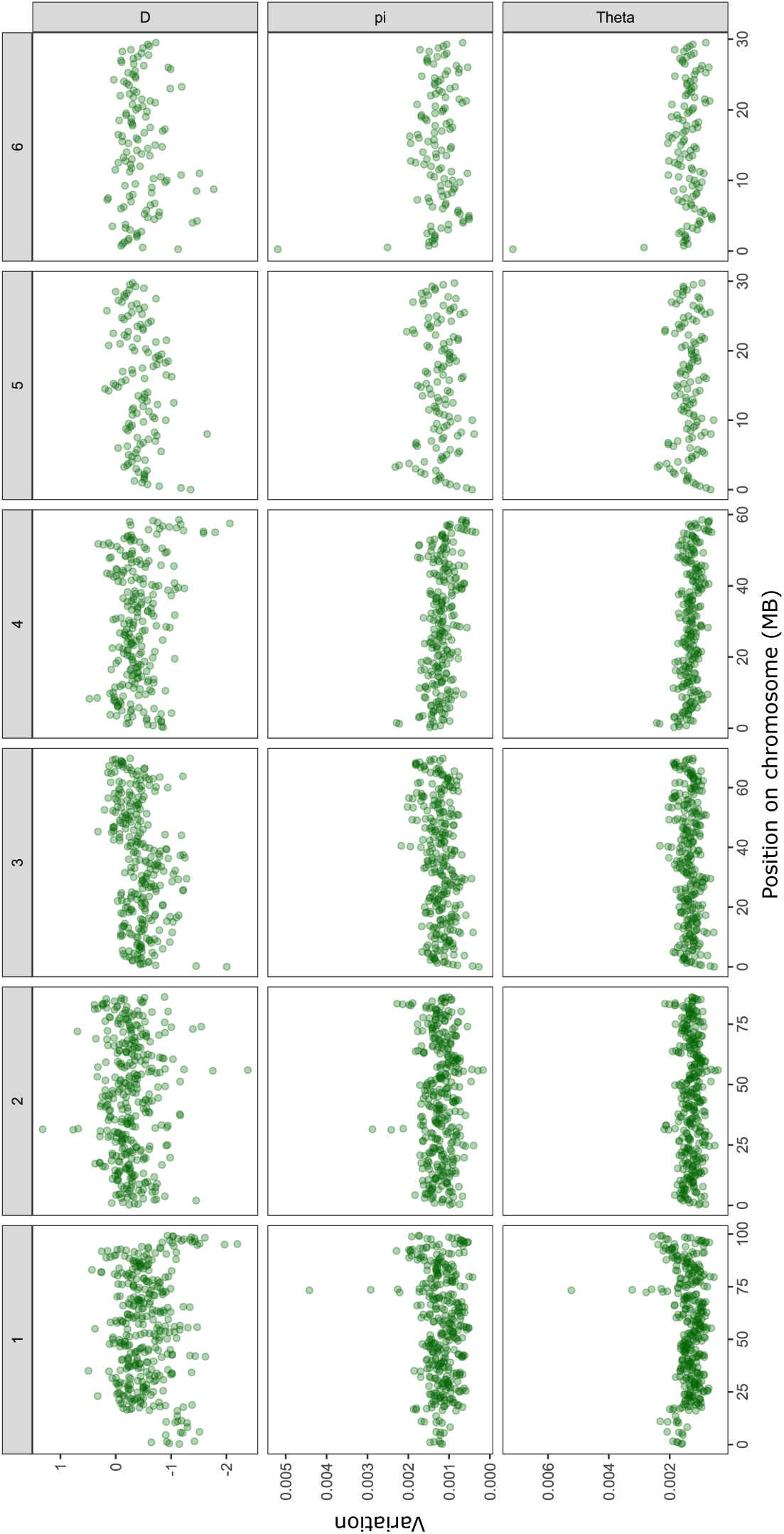
Tajima’s D, pi, and Theta for the 15 whole genome sequenced *Myzus persicae* lines. For details on aphid lines see. **Table 2**.

